# Autophagy activity is inhibited by hnRNP R

**DOI:** 10.1101/2022.06.20.496848

**Authors:** Changhe Ji

## Abstract

Autophagy is a self-eating intracellular degradation process in eukaryotic cell. Muti-pathways have been found can regulate autophagy activity through different mechanisms. In this study, we found that a nuclear abundant RNA binding protein, hnRNP R is involved in autophagy process by binding with ATG3, p62, Lamp1, LC3 and ATG9. On the other hand, hnRNP R also can regulate autophagy, we found depletion hnRNP R can activate the autophagy flux and activity. Furthermore, we also checked that autophagy has not connection with stress granules. This finding highlights some novel and nuclear located RNA binding proteins play important role in the regulation of autophagy activity.

## Introduction

Autophagy is also called macro autophagy[1-3], which is a lysosome mediated, eukaryotic cell conserved cytoplasmic degradation system supplies a protective function for the cells, and is important for cell adaptation with stressful conditions, including organelle damage, protein aggregation, nutrient deprivation, bacterial and virus invasion so on[4-6]. Recently findings implied that autophagy plays important role in Huntingtin development[7], differentiation[8], innate and adaptive immunity[9], tumour[10-12], aging[13, 14] and neuron degenerative disease, such as Alzheimer’s Disease[15], Parkinson’s Disease[16], Huntingtin Disease[17], Amyotrophic lateral sclerosis[18] and Spinal Muscular Atrophy[19].

The first step of autophagy is forming phagophore, which is an isolation membrane and derived from endoplasmic reticulum (ER) and/or the trans-Golgi and endosomes[20], Phagophore expands to engulf intra-cellular cargo, such as protein aggregates, organelles, and ribosomes, thereby sequestering the cargo in a double-membraned autophagosome[21]. The next step is lysosome will fuse matured autophagosome, and lysosomal acid proteases will mix up with autophagosome, in the end the contents in the autophagosome will be degraded. Amino acids and other by-products of degradation will be released into the cytoplasm by Lysosomal permeases and transporters[21]. In summary, the function of autophagy is removing non-functional proteins and organelles which can be considered as a cellular ‘recycling factory’ that also promotes energy efficiency through ATP generation and mediates damage control. Briefly, the procedure of autophagy is (a) phagophore formation or nucleation; (b) Atg5–Atg12 conjugation, interaction with Atg16L and multimerization at the phagophore; (c) LC3 processing and insertion into the extending phagophore membrane; (d) capture of random or selective targets for degradation; and (e) fusion of the autophagosome with the lysosome, followed by proteolytic degradation by lysosomal proteases of engulfed molecules[4].

There are many pathways to regulates autophagy,1) Nutrient Signaling. Autophagosome formation is dramatically induced when nutrient deprivation, the TOR and Ras-cAMP-PKA pathways negatively regulating autophagy will be activated[22]. 2) Insulin/Growth Factor Pathways. milieu, despite sufficient nutrients, autophagy is induced and is indispensable for maintaining cellular functions and energy production when growth factors are withdrawn from the extracellular [23]. 3) Energy Sensing. During periods of intracellular metabolic stress, activation of autophagy is essential for cell viability[22]. 4) Stress Response, such as ER stress, Hypoxia, Oxidative stress can activate autophagy. 5) Pathogen Infection, such as bacterial and virus can activate autophagy[24, 25]. 6) Transcriptional and Epigenetic Regulation of Autophagy, such as transcription activation and Chromosome Modification can affect autophagy[26]. 7) Post-Translational modification, such as Phosphorylation and Acetylation can also affect autophagy[27].

The composition of autophagy is mostly investigated, but how autophagy activity was regulated by RNA bind proteins still not clear yet so far. In this study, we found that depletion hnRNP R can activate the autophagy flux and activity. This finding highlights some novel and nuclear located RNA binding proteins play important role in the regulation of autophagy activity.

## Results

### Loss of hnRNP R activates autophagy activity

In order to investigate the role of hnRNP R in autophagy, we first knockdown long and short isoform hnRNP R by using lentivirus. Meanwhile we treat control cells and hnRNP R knockdown cells with or without autophagy inhibitor Bafilomycin A1 and we checked the proteins which are associated with autophagy by western, such as ATG9, LAMP1, p62, ATG3, LC-3 I, LC-3 II. Very interestingly, we found that ATG9 was slightly reduced in control cells but not in hnRNP R knockdown cells after Bafilomycin A1 treatment(Figure 1), ATG9 as a transmembrane protein which can protect the plasma membrane from programmed and incidental permeabilization[28],ATG9 also has the function of regulating autophagosome progression from the endoplasmic reticulum[29]. We also found hnRNP R binds with ATG9 depends on RNA (Figure 1), one explanation would be hnRNP R has some function for destabilizing ATG9. Another interesting finding is LAMP1 protein was increased in hnRNP R knockdown cells (Figure 1), we also use confocal image to confirm the downregulation of LAMP1 in hnRNP R knockdown cells (Figure 1 and Figure EV2), and this kind of increasing is not disturbed by Bafilomycin A1. LAMP1 as a Lysosome marker which plays important role in maintaining lysosomal integrity, pH and catabolism[30]. One explanation would be LAMP1 was stabilized in hnRNP R depletion cells, another possibility would be depletion hnRNP R facilities LAMP1 transcription or translation. To find out the mechanisms, more experiments need to be checked further. Next, I checked stress granule marker G3BP1, and we did not find the protein was changed in any conditions. Furtherly we checked p62, which was encoded by gene SQSTM1, is widely known as an adaptor protein of selective autophagy to promote aggregate-prone proteins for degradation. We found p62 was downregulated in control cells treatment with Bafilomycin, this result is consistent published data[31]. We also found p62 was downregulated in hnRNP R knockdown cells (Figure 1), One explanation would be p62 was destabilized in hnRNP R depletion cells, another possibility would be depletion hnRNP R inhibit p62 transcription or translation.

**Figure 1.**
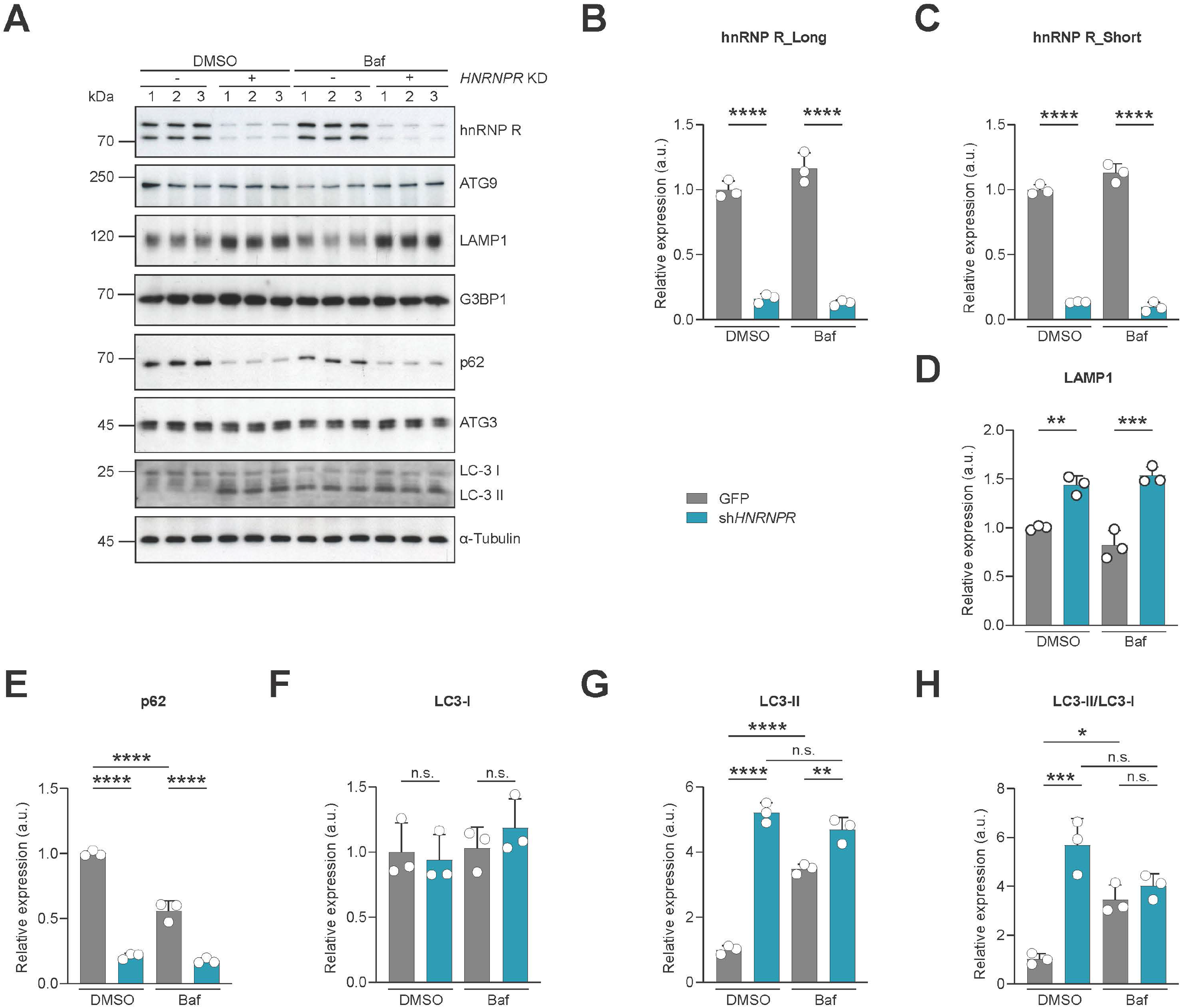
Loss of hnRNP R activates autophagy activity. A Western blot analysis of hnRNP R long, short isoform, ATG9, LAMP1, G3BP1, p62, ATG3, LC-3 I, LC-3 II and α-Tubulin protein expression in control and sh*HNRNPR* HeLa cells with or without Bafilomycin A1 treatment. B-H Quantification of hnRNP R long, short isoform, LAMP1, p62, LC-3 I, LC-3 II and LC-3 II/LC-3 I protein expression in control and sh*HNRNPR* HeLa cells with or without Bafilomycin A1 treatment after normalizing with α-Tubulin. Data are mean with SD; \**P* ≤ 0.05, \*\**P* ≤ 0.01, \*\*\**P* ≤ 0.001, \*\*\*\**P* ≤ 0.0001, n.s. not significant; unpaired two-tailed t-test (n = 3 biological replicates).

There are many ways can measure the autophagy activity in vitro or in vivo. For In vitro case, one way is using LC3-II and p62 as Indicators. LC3-II was formed after LC3 I is conjugated to PE. LC3-II is localized to both outer and inner membranes autophagosomes[32]. LC3 is currently the most widely used autophagosome marker because the amount of LC3-II reflects the number of autophagosomes and autophagy-related structures. Antibodies tend to have greater affinity for LC3-II; thus, the signal ratio of LC3-I and LC3-II does not reflect the amount ratio of cytosolic and membrane-bound LC3[33]. Furthermore, LC3-I can appear very faint depending on the antibodies and cell types. Therefore, LC3-I amount is unreliable to be used as a denominator for quantification of LC3-II (LC3-II/LC3-I). We followed the lecture’s suggestion to compare the amount of LC3-II with housekeeping proteins α-Tubulin for quantification[34]. Degradation of p62 is another widely used marker to monitor autophagic activity because p62 directly binds to LC3 and is selectively degraded by autophagy[35, 36].we found that increased LC3-II and decreased p62 in hnRNP R knockdown cells (Figure 1), this mean that autophagy activity was activated upon hnRNP R loss.

### Loss of hnRNP R activates autophagy flux

mRFP-GFP-LC3 Tandem Fluorescent Protein Quenching Assay is another very useful alternative way to measure the autophagy flux[37-40]. The reporter construct mRFP-EGFP-LC3 and the behavior of the encoded protein will be different under different pH conditions. Under neutral pH conditions, both EGFP and RFP fluorescence is observed. Under acidic pH Conditions, EGFP fluorescence is quenched, and only red fluorescence is observed. Unlipidated mRFP-EGFP-LC3 remains in the cytoplasm (light yellow) whereas lipidated mRFP-EGFP-LC3 is recruited to both inner and outer membranes of phagophores and double-membrane autophagosomes. During these steps of autophagosome formation, the fluorescent signal of both fluorophores, mRFP and EGFP, is visible and vesicles appear as yellow puncta. Under these acidic conditions, the contents within the inner membrane are eventually degraded. The green, fluorescent signal from EGFP is quenched in the acidic lysosomal conditions whereas the mRFP signal remains, resulting in red autolysosomes. The combination of green and red fluorescent signals from unlipidated mRFP-EGFP-LC3 results in a yellow background in the cytoplasm of the cells. The intensity of this yellow may change dependent upon changes in the autophagy flux. Under low autophagy conditions, most of mRFP-EGFP-LC3 remains unlipidated resulting in a yellow background and only a few yellow or red vesicles (autophagosomes and autolysosomes) are seen. After autophagy induction, many new autophagosomes form and are labelled with lipidated LC3. These rapidly fuse with lysosomes. This can be observed as an increase in the number of total vesicles and the ratio of red: yellow vesicles as well as reduced yellow background[41]. We found that depletion hnRNP R somehow can activates autophagy flux upon hnRNP R Loss (Figure 2 and Figure EV3),

**Figure 2.**
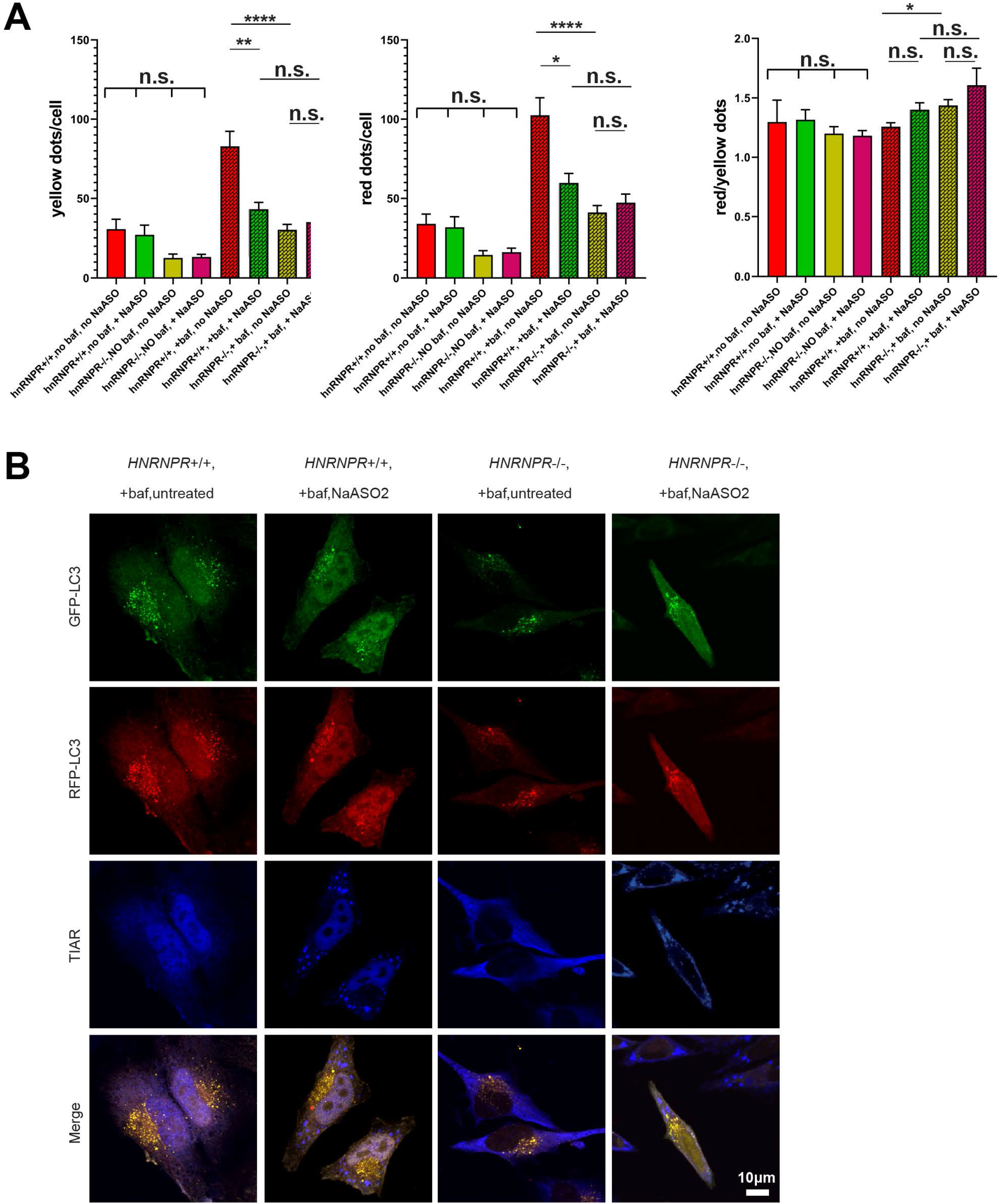
Loss of hnRNP R activates autophagy flux. A Quantification of the median number of yellow dots, red dots and red/ yellow dots in no Baf-no NaASO_2_, no Baf-with NaASO_2_, with Baf-no NaASO_2_, with Baf-with NaASO_2_ *HNRNPR*+/+ and -/- cells per cell. Data are mean with SD; \**P* ≤ 0.05, \*\**P* ≤ 0.01, \*\*\*\**P* ≤ 0.0001, n.s. not significant; unpaired two-tailed t-test (n = 3 biological replicates). B Immunostaining with antibodies against GFP-LC3, RFP-LC3 and TIAR in no Baf-no NaASO_2_, no Baf-with NaASO_2_, with Baf-no NaASO_2_, with Baf-with NaASO_2_ *HNRNPR*+/+ and -/- cells. Scale bar: 10 μm

### Loss of hnRNP R increases ubiquitin

Ubiquitin-proteasome and autophagy-lysosome systems are main pathway which clean proteins, RNAs, organelles and other toxic substrates[42].There are some functional connection between those two systems, such as when autophagy-lysosome systems was activated, Ubiquitin-proteasome also will be activated. Based on our western and confocal image results, we found that increased ubiquitin in hnRNP R depletion cells Loss (Figure 3), One possibility would be when Ubiquitin-proteasome systems activity was increased, it will generate lot of products from the UPS, to keep the cells metabolisms balance, meanwhile those of products need to be digested by autophagy-lysosome systems.

**Figure 3.**
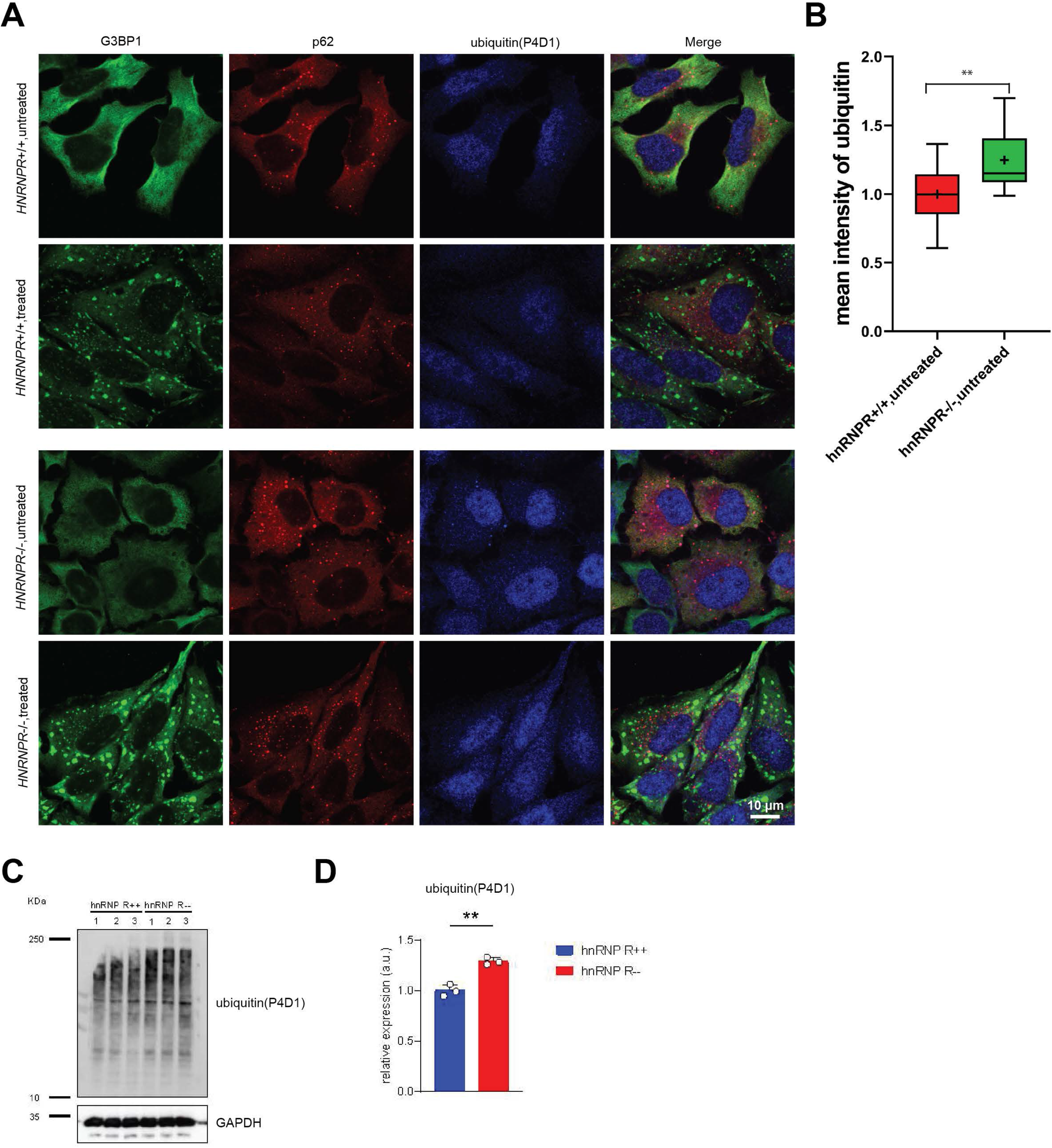
Loss of hnRNP R increases ubiquitin. A Immunostaining with antibodies against G3BP1, p62 and ubiquitin(P4D1) in without or with NaASO_2_ treatment *HNRNPR*+/+ and -/- cells. Scale bar: 10 μm B Quantification of mean intensity of ubiquitin(P4D1) in without NaASO_2_ treatment *HNRNPR*+/+ and -/- cells. Data are mean with SD; \*\**P* ≤ 0.01; unpaired two-tailed t-test (n = 3 biological replicates). C Western blot analysis of ubiquitin(P4D1) in *HNRNPR*+/+ and -/- cells. D Quantification of western band of ubiquitin(P4D1) in *HNRNPR*+/+ and -/- cells. Data are mean with SD; \*\**P* ≤ 0.01; unpaired two-tailed t-test (n = 3 biological replicates).

### Loss of hnRNP R abolishing G3BP1 binding with ATG9

Next, we want to investigate whether the connection between stress granule proteins and autophagy proteins was changed after depletion hnRNP R. we selected the core stress granule protein G3BP11 as bait to do the immunoprecipitation to check the autophagy associated protein binding[43, 44]. We found that G3BP1 binding with ATG9 was abolished upon hnRNP R depletion. Meanwhile we found that G3BP1 binds with hnRNP R and ATG9 is RNA depended (Figure 4), this would indicate that hnRNP R as an indicator to connect the G3BP1 to ATG9. G3BP1 as a substrate protein of ATG9 can be transported to the autophagosome by ATG9 and is digested by autophagy. Without hnRNP R, the stress granules proteins are more difficult to be digested, in the end has bigger stress granule formed[45].

**Figure 4.**
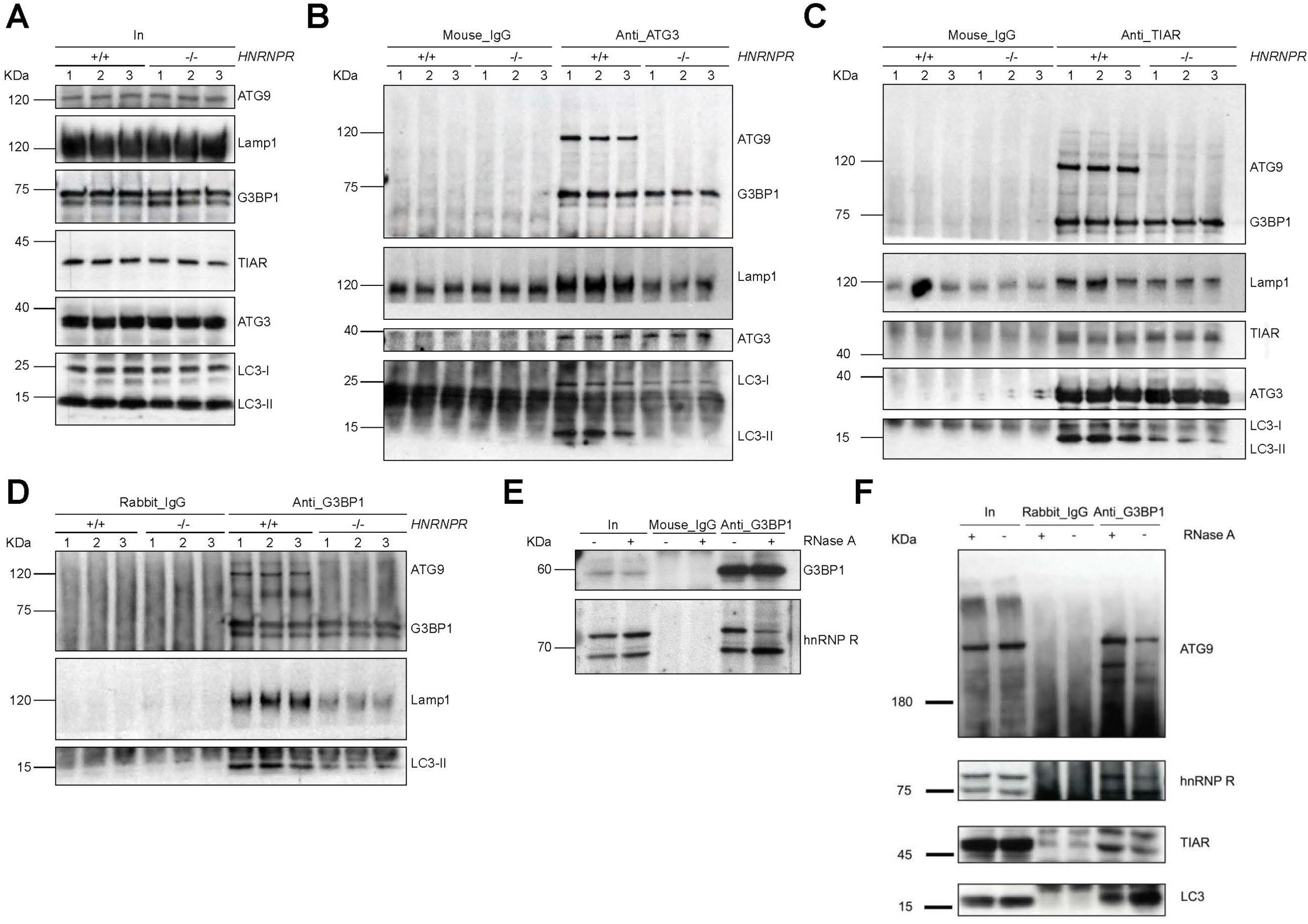
Loss of hnRNP R abolishing G3BP1 binding with ATG9. A Western blot analysis of the indicated proteins in the input (In) lysates used for co-immunoprecipitation. B Western blot analysis of ATG9, G3BP1, LAMP1, ATG3, LC3-I and LC3-II co-immunoprecipitated by an anti-ATG3 antibody. Immunoprecipitation with mouse-IgG antibody was used as control. C Western blot analysis of ATG9, G3BP1, LAMP1, TIAR, ATG3, LC3-I and LC3-II co-immunoprecipitated by an anti-TIAR antibody. Immunoprecipitation with mouse-IgG antibody was used as control. D Western blot analysis of ATG9, G3BP1, LAMP1 and LC3-II co-immunoprecipitated by an anti-G3BP1 antibody. Immunoprecipitation with rabbit-IgG antibody was used as control. E Western blot analysis of G3BP1 and hnRNP R co-immunoprecipitated by an anti-G3BP1 antibody with or without RNase A treatment. Immunoprecipitation with mouse-IgG antibody was used as control. F Western blot analysis of ATG9, hnRNP R, TIAR and LC3-I co-immunoprecipitated by an anti-G3BP1 antibody with or without RNase A treatment. Immunoprecipitation with rabbit-IgG antibody was used as control.

### Loss of hnRNP R promoting Stress granules colocalization with LAMP1

In that, G3BP1 does not bind with ATG9 after depletion hnRNP R. Next, we want to ask whether it affects the stress granule colocalization with autophagy proteins after depletion hnRNP R. We found that more stress granules colocalized with lysosome marker LAMP1 after depletion hnRNP R depended (Figure 5). Meanwhile we also found some p62 puncta also colocalized with stress granules in hnRNP R Knockout cells (Figure EV4). This is not surprised, because p62 was recruited in the autophagosome at the early stage of autophagy. One explanation would be abnormal stress granules more easily formed in hnRNP R depletion cells, meanwhile lysosome keeps closer to the abnormal stress granules and more easily to clean them. Meanwhile we also checked the stress granule positive cells which are pre-treated with Bafilomycin A1, then treatment with and wash out sodium arsenite we found that the stress granule in the cells which are treated with Bafilomycin A1 resolves faster after remove sodium arsenite for 2 hours (Figure EV5). One possibility would be lot of proteins and RNA which need to be digested were accumulation in the cytosol after Bafilomycin A1 treatment. In the end the stress granules need resolve faster to compensate consume for removing the delay digested proteins and RNA in the lysosome.

**Figure 5.**
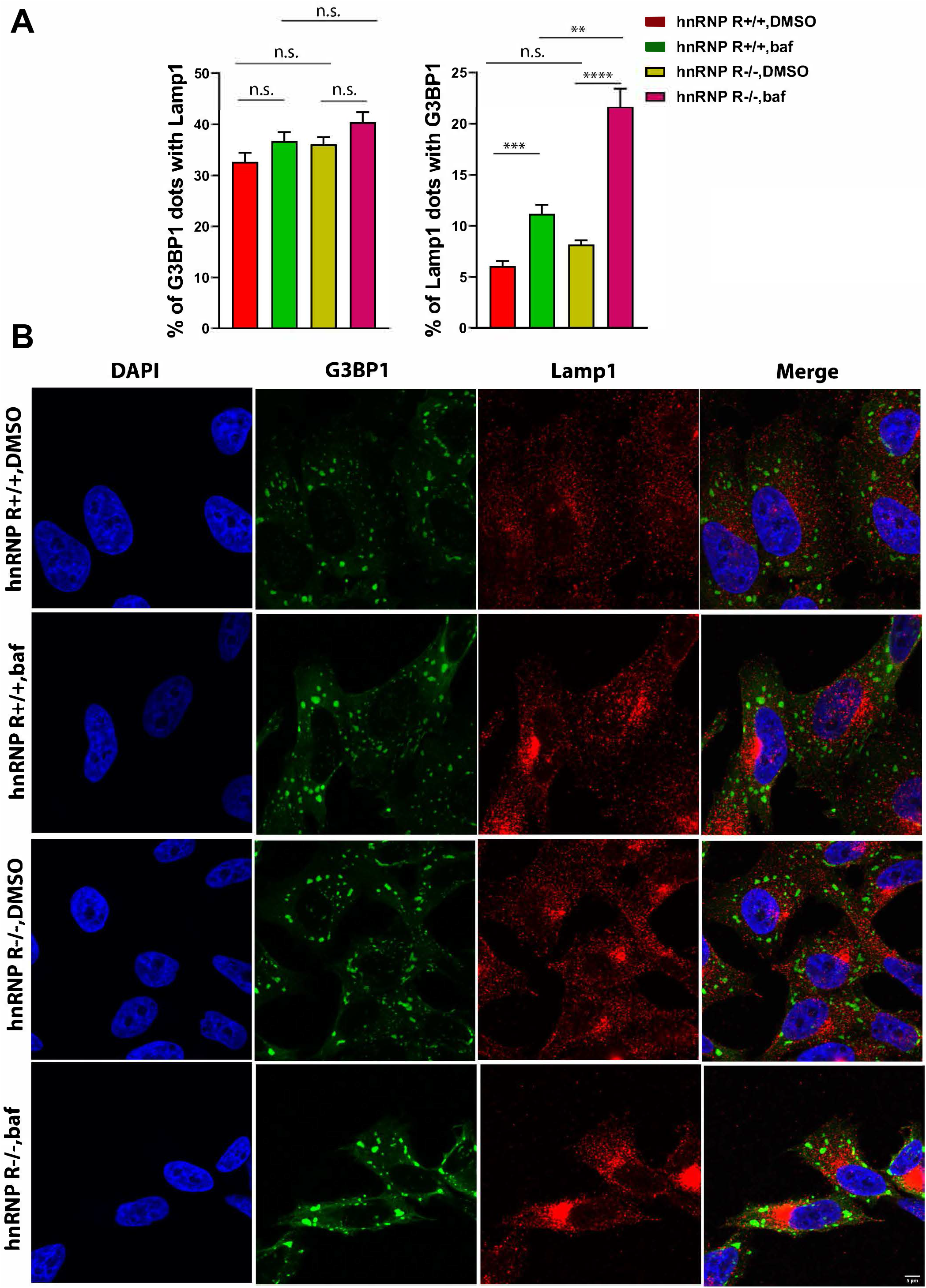
Loss of hnRNP R promoting Stress granules colocalization with LAMP1. A Quantification of the percent of stress granules colocalized with LAMP1 and the percent of LAMP1 colocalized with stress granules in *HNRNPR* +/+ and -/- cells with or without baf treatment. Data are mean with SD; \*\**P* ≤ 0.01, \*\*\**P* ≤ 0.001, \*\*\*\**P* ≤ 0.0001, n.s. not significant; unpaired two-tailed t-test (n = 3 biological replicates). B Immunostaining with antibodies against G3BP1 and LAMP1 in without or with Baf treatment *HNRNPR*+/+ and -/- cells. Scale bar: 5 μm

## Discussion

hnRNP R is a heterogeneous nuclear ribonucleoprotein, which was found mostly in the nuclear and has important functions on transcription, splicing and mRNA transportation[46-49]. Besides that, hnRNP R can also be found in the cytosol and motoneuron axons[47, 48, 50], which plays important role in mRNA translation, mRNP transport and stress granule formation[51]. In this study, we found that hnRNP R also plays important role in regulation of autophagy activity in the cytosol. We used hnNRP R knockdown and knockout cells both found that depletion hnRNP R can promote LC3-II formation. Meanwhile we used mRFP-GFP-LC3 Tandem Fluorescent Protein Quenching Assay proving that depletion hnRNP R increase autophagy activity. We also prove that depletion hnRNP R activate ubiquitin-proteasome system. Furtherly, we found loss of hnRNP R abolishing G3BP1 binding with ATG9 and promoting stress granules colocalization with LAMP1. We propose that, Depletion hnRNP R will have a series of effects for the cells from nuclear to the cytosol. Some pathways need to be activated or inactivated for the adaption.

One thing must note that, we found acute knockdown hnRNP R by using lentivirus or chronic knockout hnRNP R by using prime editing has different phenotype for the autophagy. p62 was reduced in hnRNP R knockdown cells but a little bit decreases in hnRNP R knockout cells. One possibility would be acute hnRNP R knockdown same as starvation, need rapid nutrition consumption to keep the cells survive[52]. However, as mentioned in hnRNP R knockout cells, after a long period of adaptation, some other pathways are adjusted to make the p62 tend to stable (Figure EV1).

Besides p62, another obvious difference is LC3 in knockdown or knockout cells. You will find LC3-II is increased in hnRNP R knockdown cells but reduced in hnRNP R knockout cells in basic condition. One explanation would be acute hnRNP R knockdown same as starvation, need rapid digestion to keep the cells survive. However, after long period of adaptation, some other pathways are adjusted to brief shot down autophagy to save nutrition in hnRNP R knockout. Very interestingly, LC3-II both are increased in hnRNP R knockdown cells and knockout cells after Bafilomycin A1 treatment. This not surprise, high concentration of Bafilomycin A1 as a highly specific inhibitor for autophagy and is toxic for the cells. Cells need activate autophagy as much as possible to keep alive (Figure EV1).

hnRNP R affecting G3BP1 binding with ATG9 is another surprise finding. G3BP1 has been found extensively connect with autophagy proteins. One research found that G3BP1 act in a non-redundant manner to anchor the tuberous sclerosis complex (TSC) protein complex to lysosomes and suppress activation of mTORC1[53]. ATG9 has important function in early stage of autophagosome formation[54], Consistent with previous finding by other research[55], we also found that G3BP1 can also bind with ATG9, this indicates ATG9 can also tether the stress granule protein to the autophagosome, we also found that hnRNP R link the binding of G3BP1 with ATG9. One possibility is G3BP1 first binds with hnRNP R, then binds with ATG9. Another possibility would be G3BP1 with other proteins forming the complex and depletion hnRNP R changed the conformation of the complex, in the end the G3BP1 cannot bind with ATG9.

Disturbances in autophagy and stress granule dynamics have been implicated as potential mechanisms underlying neuronal disorders. Yet the relationship between autophagy and stress granule dynamics remains poorly characterized. Previous we found that depletion hnRNP R stimulates stress granule formation, alters stress granules phenotype, and impair stress granules resolving. In this study we also found loss of hnRNP R promoting stress granules colocalization with LAMP1. This closer connection between stress granule and lysosome indicates that autophagy has higher activity for stress granule clearance after depletion hnRNP R. In summary, in the study, this highlights hnRNP R as an RNA binding protein has important function for autophagy activity regulation. It provides a valuable reference to the autophagy dysfunction related disease, such as neuron degenerative disease.

## Materials and Methods

### HeLa cell culture

HeLa cells (Leibniz Institute DSMZ-German Collection of Microorganisms and Cell Cultures GmbH, DSMZ no. ACC 57) were cultured at 37°C and 5% CO_2_ in high glucose Dulbecco’s Modified Eagle Medium with GlutaMAX™ Supplement (DMEM; Gibco) supplemented with 1 mM sodium pyruvate (Gibco), 10% fetal calf serum (Linaris) and 1% Penicillin-Streptomycin (Gibco). Cells were passaged when they were 80-90% confluent. Cells tested negative for mycoplasma contamination.

### Preparation of lentiviral knockdown and GFP-RFP-LC3 constructs

shRNAs targeting human *HNRNPR* were cloned into a modified version of pSIH-H1 shRNA vector (System Biosciences) containing EGFP according to the manufacturer’s instructions. The following antisense sequences were used for designing shRNA oligonucleotides targeting *HNRNPR*: 5’-GTTCTGCTTCCTTGAATATGA -3’, (Supplementary Table 1). GFP-RFP-LC3 lentiviral construct was a gift from Dr. Patrick Lüningschrör (Institute of Clinical Neurobiology, University Hospital of Wuerzburg). Empty pSIH-H1 expressing EGFP was used as control. Lentiviral particles were packaged in HEK293T cells with pCMV-pRRE, pCMV-pRSV, and pCMV-pMD2G as described before[56].

### Generation of hnRNP R knockout HeLa cell lines by prime editing

Knockout HeLa cells were generated by prime editing[57]. The sgRNA sequence 5’-CAAGGTGCAAGAGTCCACAA-3’ was identified in exon 4 of *HNRNPR* using the Broad Institute sgRNA design tool (https://portals.broadinstitute.org/gpp/public/analysis-tools/sgrna-design) and a pegRNA was designed for insertion of an in-frame stop codon. The pegRNAs were cloned into pU6-pegRNA-GG-acceptor (Addgene Plasmid #132777) [57] as following. 1 µg pU6-pegRNA-GG-acceptor was digested with Fast Digest Eco31I (IIs class) (Thermo Fisher Scientific) in a 20 µl reaction. Following agarose gel electrophoresis, the vector backbone was gel-purified using the NucleoSpin Gel and PCR Clean-up kit (Macherey-Nagel). Oligonucleotides pegRNA3-1 and pegRNA3-2 (Appendix Tables S1 and S2) were annealed and extended in a 30 µl PCR reaction containing 0.4 µM each of pegRNA3-1 and pegRNA3-2, 3 µl of 10× Extra Buffer, 0.33 mM of each dNTP and 1 U of Taq DNA polymerase (VWR) using one cycle of 94°C for 5 min 30 s, 60°C for 30 s and 72°C for 30 s. The pegRNA was assembled and inserted into pU6-pegRNA-GG-acceptor in a 10 µl reaction containing 25 ng digested pU6-pegRNA-GG-acceptor, 1.125 µl PCR reaction from the previous step, 45 nM pegRNA3-3AGTGA (Appendix Table S1) and 5 µl NEBuilder HiFi DNA Assembly Master Mix (NEB) and incubated at 37°C for 1 h.

For transfection, 10^5^ HeLa cells were seeded in a 12-well plate in a volume of 0.5 ml of DMEM 24 h prior transfection. After 24 h, cells were co-transfected with 750 ng of pCMV-PE2 Plasmid (Addgene Plasmid #132775) and 250 ng of assembled pU6-pegRNA-GG-acceptor using Lipofectamine 2000 (Invitrogen). Cells were harvested 72 h after transfection by trypsinization using TrypLE™ Express Enzyme (Gibco), counted and diluted count to 9 cells per ml DMEM medium. 100 µl of cell suspension were added per well of a 96-well plate. Cells were grown for 7-10 d to allow formation of colonies from single cells. Colonies were trypsinized, transferred to a 48-well plate and grown for another 7 d for cell expansion. Then, cells in each well were trypsinized and split into two wells of a 24-well plate. One of these wells was used for genotyping with primers listed in Appendix Tables S2.

### Bafilomycin A1 and sodium arsenite treatment

HeLa cells were grown on 10 mm glass coverslips for 2 d. Add Bafilomycin A1 (Cell Signaling Technology) in the medium with final concentration is 1 μM or DMSO as control and incubated for 4 hours prior to harvesting. Add sodium arsenite (sigma) in the medium with final concentration is 0.5 mM or water as control and incubated for 1 hour prior to harvesting.

### Immunofluorescence staining

For immunofluorescence staining, HeLa cells were grown on 10 mm glass coverslips for 2 d. Cells were washed three times with pre-warmed phosphate-buffered saline (PBS), fixed with 4% paraformaldehyde (PFA; Thermo Fisher Scientific) for 20 min at room temperature followed by three washes with pre-warmed PBS. For permeabilization, 0.3% Triton X-100 was applied for 20 min at room temperature followed by three washes with pre-warmed PBS. For reduction of unspecific binding, cells were treated with 10% horse serum and 2% bovine serum albumin (BSA) in Tris-buffered saline with Tween 20 (TBS-T) for 0.5 h at room temperature. Primary antibody was applied overnight at 4°C followed by three washes with TBS-T and incubation with fluorescently labeled secondary antibodies in TBS-T for 1 h at room temperature. Cells were washed three times with TBS-T at room temperature, incubated with DAPI (Sigma-Aldrich) diluted 1:1,000 in PBS for 10 min and washed once with water. Cells were embedded with Aqua-Poly/Mount (Polysciences). The primary and secondary antibodies used for immunostaining are listed in Appendix Table S3.

### Confocal microscopy

Images were acquired on an Olympus Fluoview 1000 confocal system equipped with four excitation lasers: a 405nm, a 473nm, a 559nm and a 635nm laser. A 60× objective (oil differential interference contrast, numerical aperture: 1.35) was used for image acquisition. For quantification of immunofluorescence signals, raw images were projected using ImageJ/Fiji as average intensity and mean gray values were measured after background subtraction.

### Total protein extraction

HeLa cells were washed once with ice-cold DPBS (without MgCl_2_, CaCl_2_; Sigma-Aldrich) and collected by scraping. Cells were lysed in 1 ml lysis RIPA buffer (20 mM Tris-HCl pH 7.5, 150 mM NaCl, 1 mM Na_2_EDTA, 1 mM EGTA, 1% NP-40, 1% sodium deoxycholate, 2.5 mM sodium pyrophosphate, 1 mM β-glycerophosphate) on ice for 15 min. Lysates were centrifuged at 16,000×g for 15 min at 4 °C and the supernatant was transferred into a new tube. Protein concentration was measured by Pierce BCA Protein Assay Kit (Thermo Fisher Scientific). Lysates were dissolved in 5× Laemmli buffer (300 mM Tris-HCl pH 6.8, 10% SDS, 50% glycerol, 0.05% Bromophenol Blue, 100 mM DTT) and analyzed by Western blotting using antibodies listed in Table S3.

### Immunoprecipitation

Cells were washed once with ice-cold DPBS, collected by scraping and lysed in 1 ml lysis buffer (10 mM HEPES pH 7.0, 100 mM KCl, 5 mM MgCl_2_, 0.5% NP-40) on ice for 15 min. Lysates were centrifuged at 20,000×g for 15 min at 4°C. For co-immunoprecipitation of histone H3, lysates were sonicated prior to centrifugation. 10 µl of Dynabeads Protein G or A (depending on the antibody species) (Thermo Fisher Scientific) and 1 µg antibody or IgG control were added to 200 µl lysis buffer and rotated for 30–40 min at RT. Then 200 µl lysate was added to the antibody-bound beads and rotated for 2 h at 4°C. For RNase treatment, 3 µl RNase A (Thermo Fisher Scientific) were added to 200 µl lysate and incubated at 37°C for 10 min before proceeding with co-immunoprecipitation. Beads were washed twice with lysis buffer and proteins were eluted in 1× Laemmli buffer. Proteins were size-separated by SDS-PAGE and analyzed by Western blotting. The intensity of protein bands was measured with ImageJ/Fiji, and co-immunoprecipitation was quantified by normalization to input and to the efficiency of bait protein purification. The antibodies used for immunoprecipitation and Western blotting are listed in the Appendix Table S3.

## Supporting information

Autophagy_primer and antibody list

## Expanded View Figure legends

**Figure EV1.**
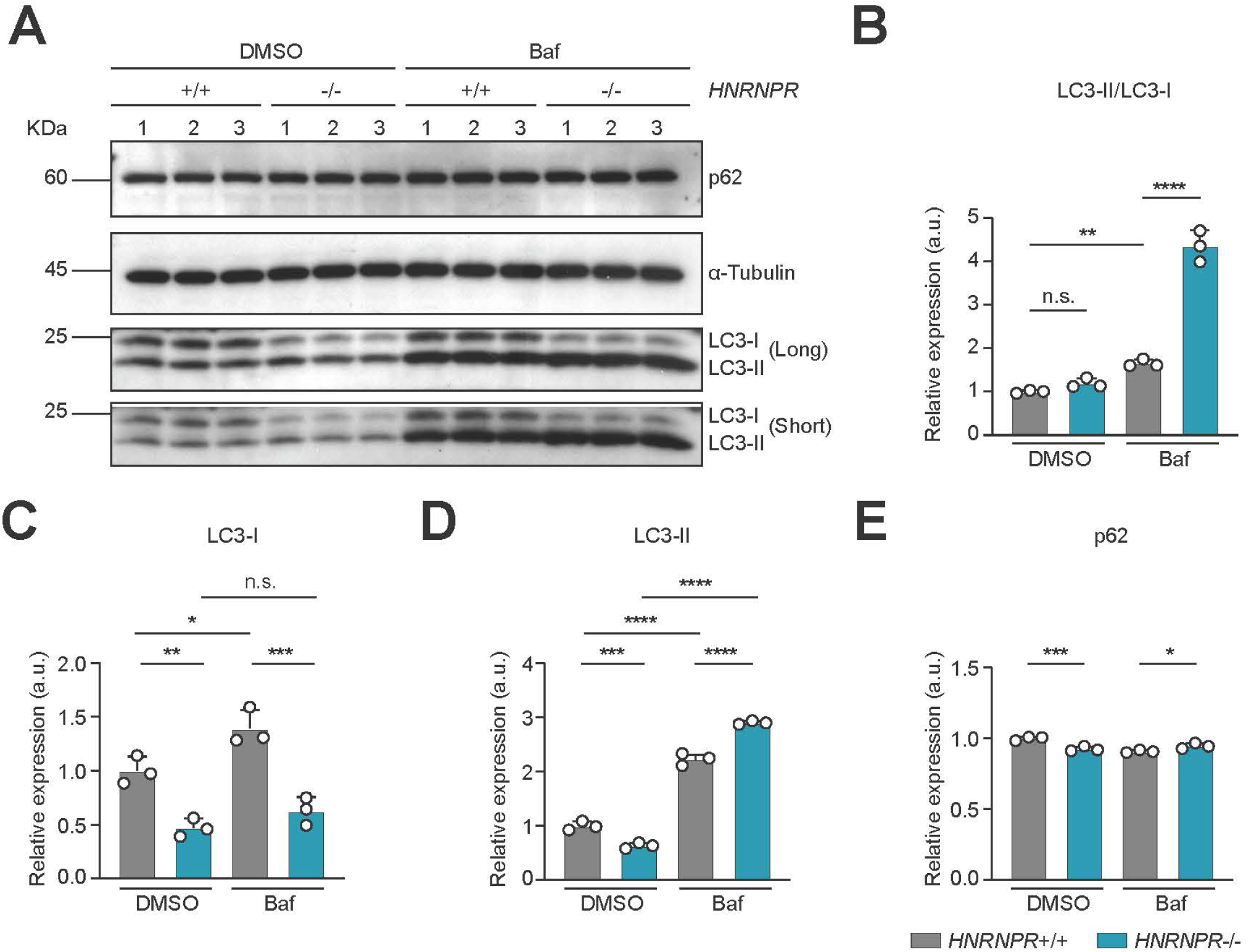
Increased autophagy activity in hnRNP R knock out cells. A Western blot analysis of p62, LC-3 I, LC-3 II and α-Tubulin protein expression in *HNRNPR*+/+ and -/- cells HeLa cells with or without Bafilomycin A1 treatment. B-E Quantification of LC-3 II/LC-3 I, LC-3 I, LC-3 II and p62 protein expression in *HNRNPR*+/+ and -/- cells HeLa cells with or without Bafilomycin A1 treatment after normalizing with α-Tubulin. Data are mean with SD; \**P* ≤ 0.05, \*\**P* ≤ 0.01, \*\*\**P* ≤ 0.001, \*\*\*\**P* ≤ 0.0001, n.s. not significant; unpaired two-tailed t-test (n = 3 biological replicates).

**Figure EV2.**
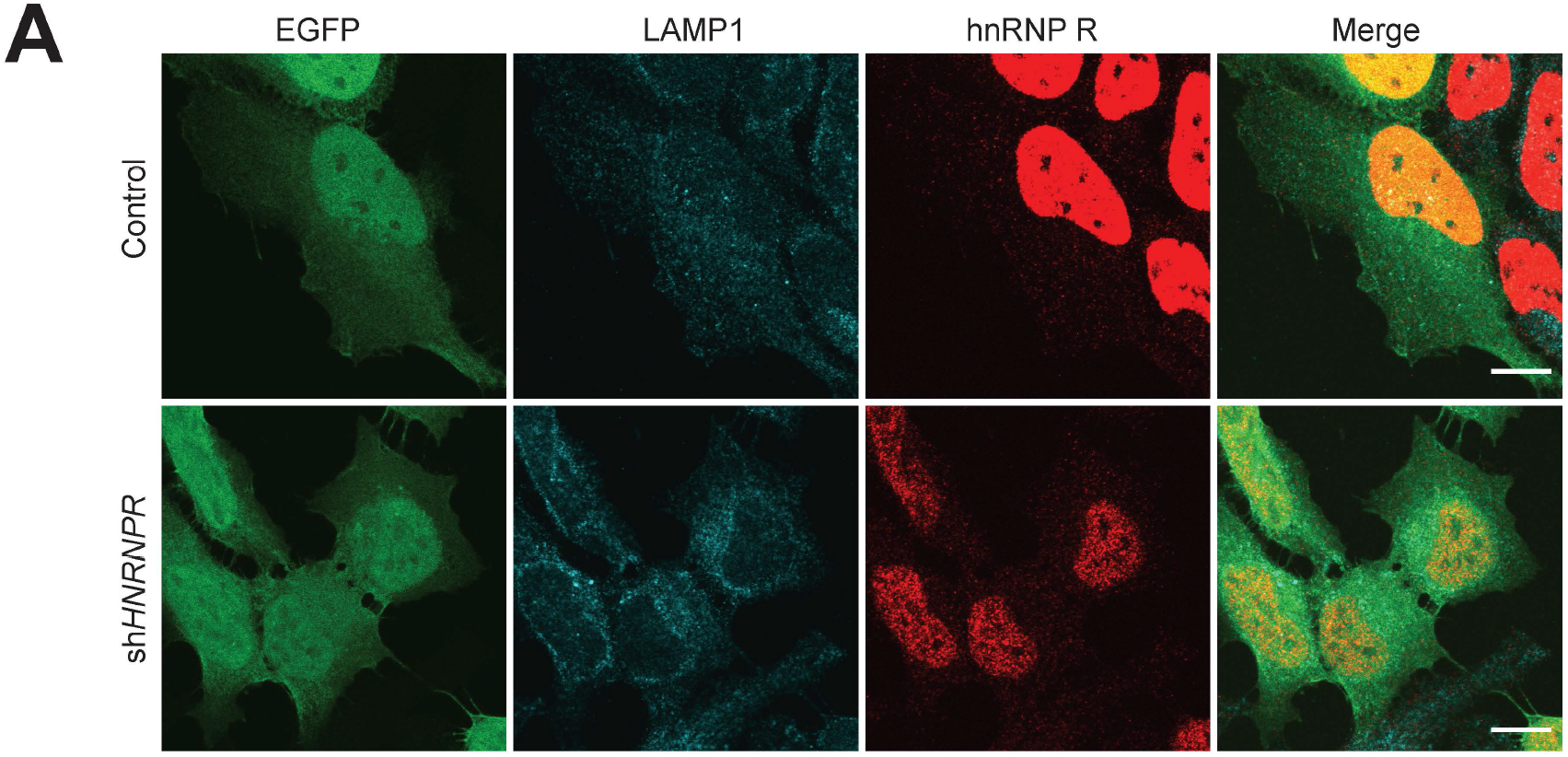
Increased LAMP1 in hnRNP R knockdown cells. A Immunostaining with antibodies against EGFP, LAMP1 and hnRNP R(Abcam) in Control and sh*HNRNPR* HeLa cells. Scale bar: 10 μm.

**Figure EV3.**
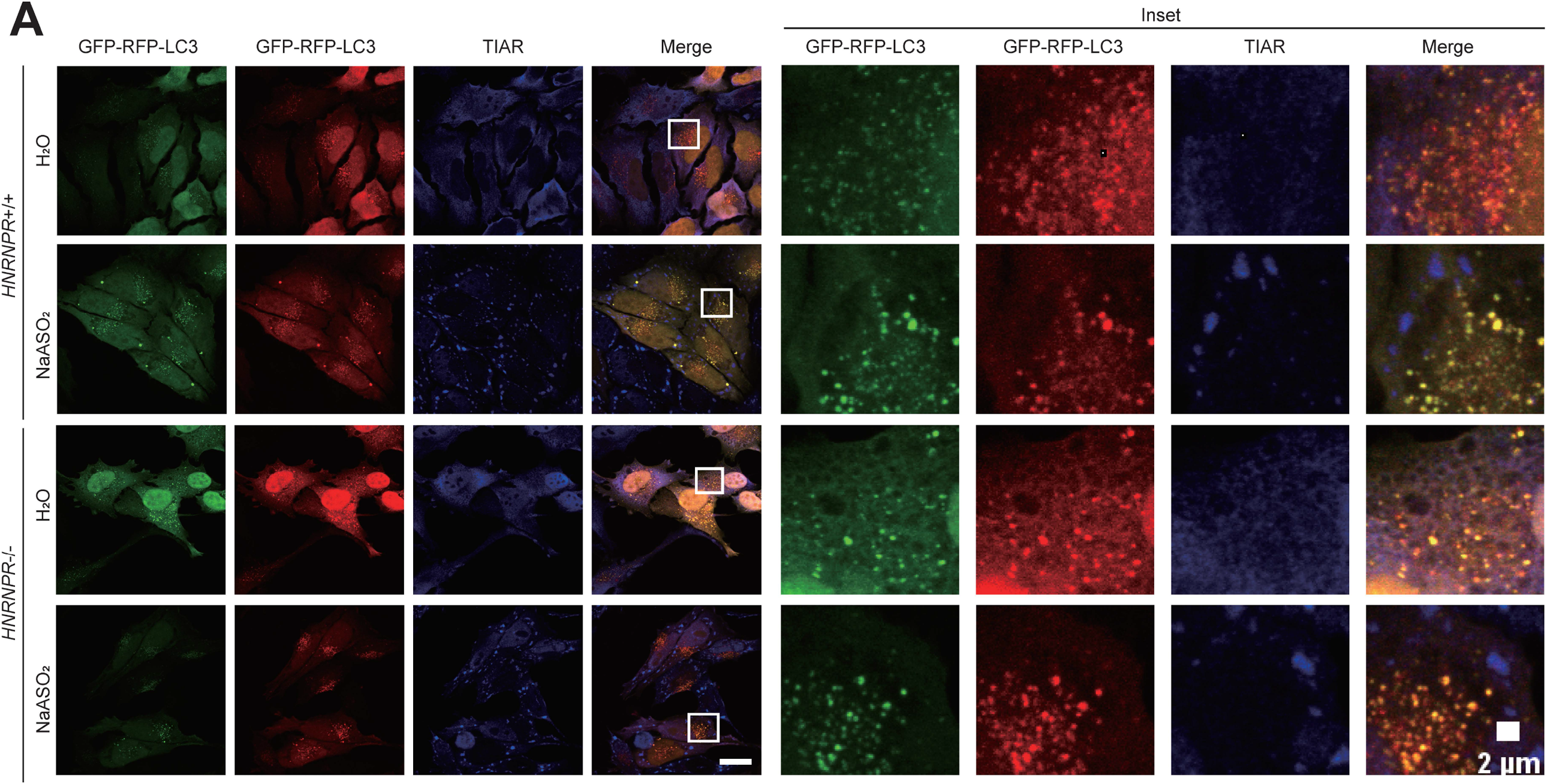
Stress granules do not colocalize with LC3 puncta. A Immunostaining with antibodies against TIAR and visualized GFP-LC3,RFP-LC3 in *HNRNPR*+/+ and -/- cells without or with NaASO_2_ treatment. Scale bar: 10 μm

**Figure EV4.**
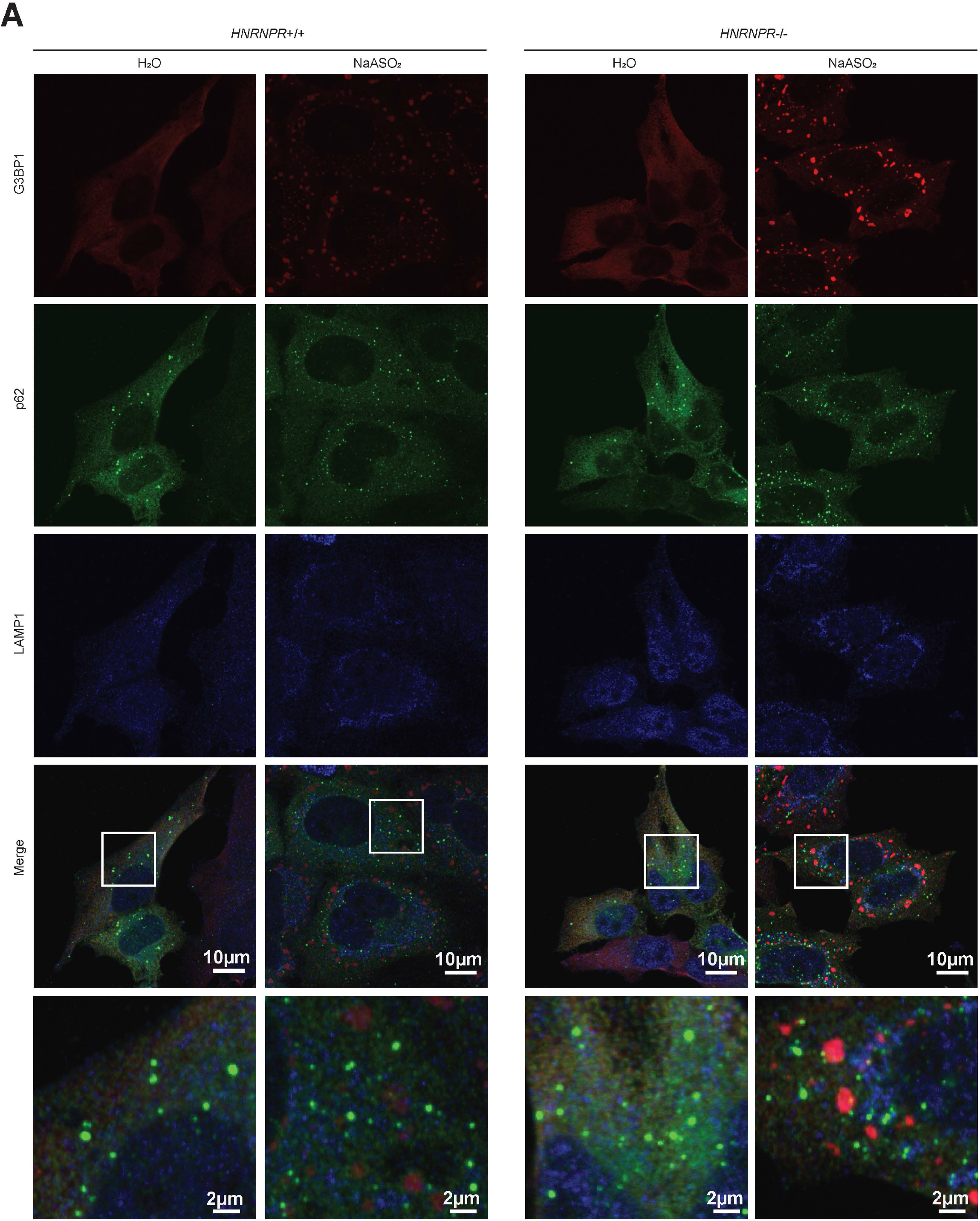
Stress granules do not colocalize with p62 puncta. A Immunostaining with antibodies against G3BP1, LAMP1 and p62 in in *HNRNPR*+/+ and -/- cells without or with NaASO_2_ treatment. Scale bar: 10 μm

**Figure EV5.**
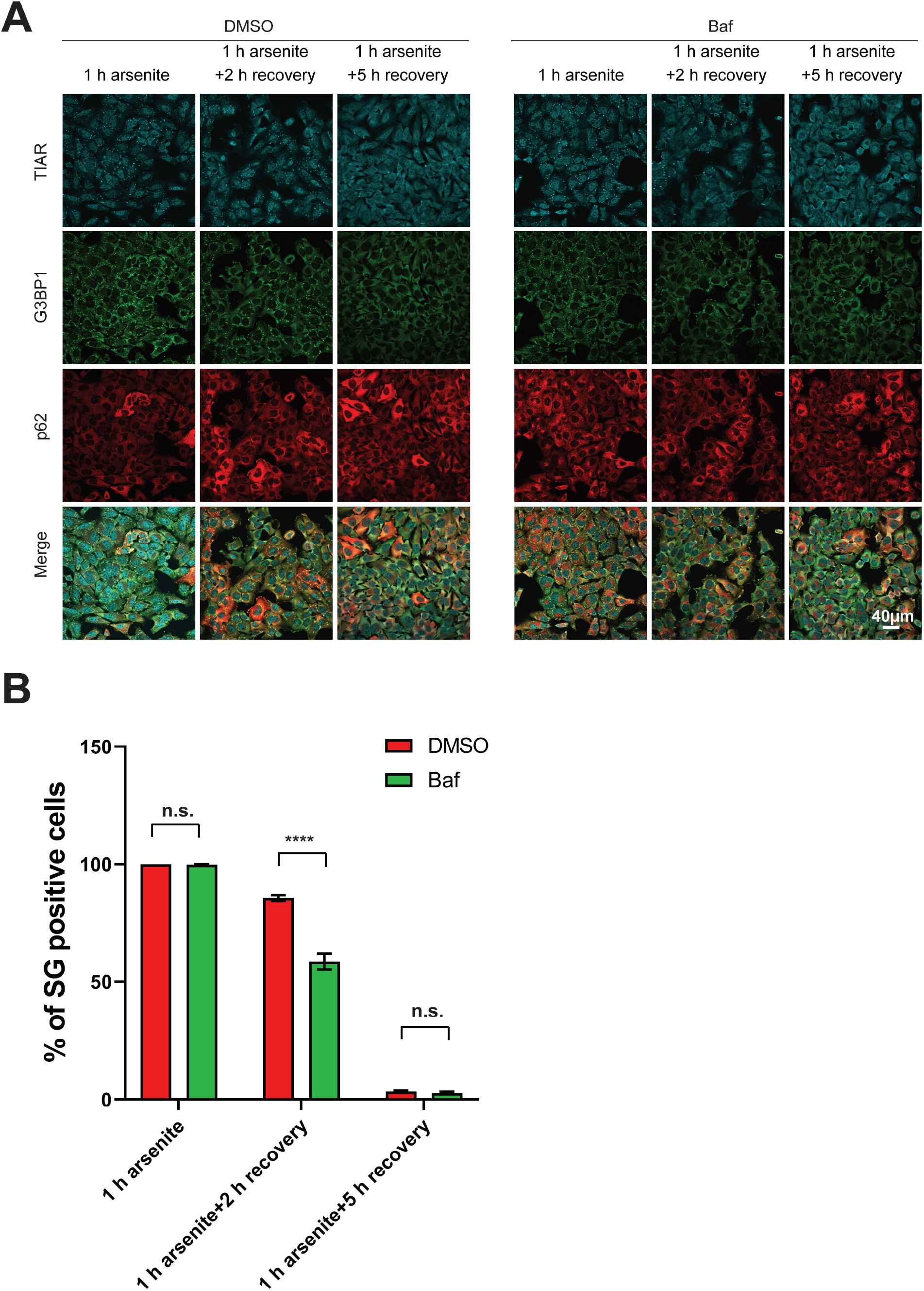
Bafilomycin A1 facilitates Stress granules dissolving. A Immunostaining with antibodies against TIAR, G3BP1 and p62 from HeLa cells which are pre-treated without or with Bafilomycin A1 for four hours, then treated with NaASO_2_ for one hour, then removed NaASO_2_ with two or five hours. Scale bar: 40 μm B Quantification of the percent of stress granules positive cells which are pre-treated without or with Bafilomycin A1 for four hours, then treated with NaASO_2_ for one hour, then removed NaASO_2_ with two or five hours. Data are mean with SD; \*\*\*\**P* ≤ 0.0001, .s. not significant; unpaired two-tailed t-test (n = 3 biological replicates).

