## Supplementary material for "Autophagy activity is inhibited by hnRNP R": Autophagy_primer and antibody list

### hnRNP R is a regulator of stress granule formation

Changhe Ji

**Table of Contents:**

Appendix Table S1 - Sequences of oligonucleotides for shRNA cloning

Appendix Table S2 - Sequences of oligonucleotides for *HNRNPR* prime editing

Appendix Table S3 - Antibodies

#### SUPPLEMENTARY TABLES

##### Supplementary Table 1: Sequences of oligonucleotides for shRNA cloning

| Name | Sequence (5'-3') |
|---|---|
| Sh*HNRNPR*_F | GAT CCG TTC TGC TTC CTT GAA TAT GAT CAA GAG TCA TAT TCA AGG AAG CAG AAC TTT TTG |
| Sh*HNRNPR*_R | AAT TCA AAA AGT TCT GCT TCC TTG AAT ATG ACT CTT GAT CAT ATT CAA GGA AGC AGA ACG |

##### Supplementary Table 2: Sequences of oligonucleotides for *HNRNPR* prime editing

| Name | Sequence (5'-3') |
|---|---|
| pegRNA3-1 | TAT CTT GTG GAA AGG ACG AAA CAC CGC AAG GTG CAA GAG TCC ACA AGT TTT AGA GCT AGA AAT AGC |
| pegRNA3-2 | GCA CCG ACT CGG TGC CAC TTT TTC AAG TTG ATA ACG GAC TAG CCT TAT TTT AAC TTG CTA TTT CTA GCT CTA AAA C |
| PegRNA3-3AGTGA | TGA AAA AGT GGC ACC GAG TCG GTG CTC AGG TCC CTT TGT CAC TTG GAC TCT TGC ACC TTT TTT TAA GCT TGG GCC GCT CGA G |
| EXON4_F | TCCGACATCTGGCAAAGACA |
| EXON4_F (for AGTGA) | AGG TGC AAG AGT CCA AGT GA |
| EXON4_R | GGTCAAATGCCTCTTTCCATGT |

**Supplementary Table 3: Antibodies used for Western blotting (WB), immunoprecipitation (IP), immunofluorescence (IF)**

| Antibody | Application | Dilution | Source | Identifier |
|---|---|---|---|---|
| Rabbit monoclonal Anti-ATG9A | WB | 1:1000 | Abcam | Cat# ab108338; RRID:AB_ AB_10863880 |
| Rabbit polyclonal anti-ATG9A | WB | 1:1000 | Novus Biologicals | Cat# NB110-56893; RRID: AB_837629 |
| Mouse monoclonal anti-LAMP-1 Antibody (H4A3) | WB<br>IF | 1:2000<br>1:100 | Santa Cruz | Cat# sc-20011. RRID: AB_626853 |
| Rabbit monoclonal Anti-LAMP1 (D2D11) | WB | 1:1000 | Cell Signaling Technology | Cat# 9091S; RRID: AB_2687579 |
| Guinea pig polyclonal anti-p62/ SQSTM1 | WB<br>IF | 1:2000<br>1:100 | PROGEN Biotechnik | Cat#GP62-C; RRID: AB_1542690 |
| Rabbit polyclonal anti-hnRNP R | WB | 1:4000<br>1:200 | Abcam | Cat#ab30930; RRID:AB_2295532 |
| Rabbit polyclonal anti-hnRNP R (N-term) | WB | 1:1000 | Abgent | Cat#AP17239a; RRID: AB_11136203 |
| Rabbit polyclonal anti-hnRNP R (C- term) | IF | 1:200 | Sigma | Cat# HPA026092; RRID: AB_1850885 |
| mouse monoclonal anti-Atg3 (A-3) | WB<br>IF | 1:2000<br>1:100 | Santa Cruz | Cat# sc-393660; |
| Rabbit polyclonal anti-LC3B | WB | 1:1000 | Novus Biologicals | Cat# NB100-2220; RRID: AB_10003146 |
| Mouse monoclonal anti-α-Tubulin | WB | 1:2000 | Sigma-Aldrich | Cat# T5168 RRID: AB_477579 |
| Mouse monoclonal anti-Ubiquitin(P4D1) | WB<br>IF | 1:2000<br>1:100 | Enzo Life Sciences | Cat#BML-PW0930-0100; RRID: AB_11181462 |
| Mouse monoclonal anti-TIAR | IF<br>WB | 1:400<br>1:2000 | BD | Cat#610352; RRID:AB_397742 |
| Rabbit polyclonal anti-G3BP1 | IF | 1:500 | Proteintech | Cat# 13057-2-AP; RRID: AB_2232034 |
| Mouse monoclonal anti-hnRNP A1 (clone 4B10) | WB | 1:2000 | Santa Cruz | Cat#sc-32301; RRID: AB_627729 |
| Mouse monoclonal anti-G3BP1 | IF | 1:500 | BD | Cat# 611127; RRID:AB_398438 |
| Mouse monoclonal anti-GAPDH (clone 6C5) | WB | 1:3000 | Calbiochem | Cat# CB1001; RRID: AB_2107426 |
| Chicken polyclonal anti-GFP | IF | 1:1000 | Abcam | Cat# ab13970; RRID: AB_300798 |

| Antibody | Application | Dilution | Source | Identifier |
|---|---|---|---|---|
| Mouse IgG control | IP | | Santa Cruz | Cat#sc-2025; RRID:AB_737182 |
| Goat polyclonal anti-Mouse,Peroxidase conjugated | WB | 1:5000 | Jackson ImmunoResearch | Cat#115-035-146. RRID: AB_2307392 |
| Donkey polyclonal anti-Rabbit, Peroxidase conjugated | WB | 1:5000 | Jackson ImmunoResearch | Cat#711-035-152; RRID:AB_10015282 |
| Donkey polyclonal anti-Goat,Peroxidase conjugated | WB | 1:5000 | Jackson ImmunoResearch | Cat#705-035-003; RRID: AB_2340390 |
| Donkey polyclonal anti-Chicken, Alexa Fluor® 488 conjugated | IF | 1:800 | Jackson ImmunoResearch | Cat#703-545-155; RRID: AB_2340375 |
| Donkey polyclonal anti-Mouse, Cy™3 conjugated | IF | 1:800 | Jackson ImmunoResearch | Cat#715-165-151. RRID: AB_2315777 |
| Donkey polyclonal anti-Rabbit, Cy™5 conjugated | IF | 1:800 | Jackson ImmunoResearch | Cat#711-175-152; RRID: AB_2340607 |
